# Dynamic Micropatterning Reveals Spatial Dynamics of B Cell Receptor Signaling and Immune Synapse Formation

**DOI:** 10.1101/2025.08.11.669641

**Authors:** Blanca Tejeda-González, Sara Hernández-Pérez, Elmeri Kiviluoto, Johanna Ivaska, Pieta K. Mattila

## Abstract

B cell activation by foreign antigens, recognized by the B cell receptor (BCR), is the key prerequisite for cell differentiation for antibody production. B cells typically encounter antigens bound to the surface of antigen presenting cells (APCs), leading to the formation of the immunological synapse (IS). To gain a deeper understanding of lymphocyte activation, we developed a dynamic micropatterning technique that enables the modeling of IS formation with exceptional spatial and temporal control. Using this method, we can image B cells before and after BCR engagement, in both fixed and live samples. We compared the activation of different BCR proximal signaling proteins in activatory and non-activatory areas of the synapse and found that the activated signaling proteins exhibited distinct spatial distributions. While pCD79A was strongly localized in the antigen-tethered area, surprisingly, pPLCγ2 was enriched in regions lacking BCR ligands. We also visualized the formation of the IS in living cells using enhanced-resolution microscopy in 3D. We identified different cell behaviors during this process, including the repurposing of pre-existing actin-based protrusions as ready-made building blocks for the IS — a feature uniquely detectable with this highly controllable system. Extending our approach to include a co-stimulatory B cell ligand, ICAM-1, and T cell system using CD3 and CD28 antibodies as ligands, we demonstrate the broader applicability of this method. Overall, our results highlight the power of dynamic micropatterning in elucidating the rapid and dynamic earliest steps of the IS formation with high spatial and temporal precision.

## Introduction

B lymphocytes (B cells), as a key component of the adaptive immune system, are responsible for mounting humoral immune responses against pathogens. Recognition of specific antigens via the B cell receptor (BCR), along with additional co-stimulatory signals, initiates B cell activation and differentiation into antibody-producing plasma cells. It is increasingly recognized that, while soluble and particulate antigens can activate B cells, they typically encounter antigens in vivo that are displayed on the surface of antigen-presenting cells (APCs). This interaction leads to the formation of an intricate cell-cell contact known as the immunological synapse (IS) (*Harwood & Batista, 2010*; *Huang et al*., 2005; *Kuokkanen et al*., 2015; *Tolar*, 2017; *Wykes et al*., 1998), which shares strong similarity with the synapses originally described in T cell activation (*Grakoui et al*., 1999; *Monks et al*., 1998). The IS displays a bull’s-eye-like structure with a central supramolecular activation cluster (cSMAC) where BCR-antigen com-plexes gather, and a peripheral SMAC (pSMAC) that consists of a radial, actin-rich protrusion enriched in adhesion molecules. The formation of the IS is initiated by cell spreading over the antigen-presenting surface, enabling engagement with a high density of antigen, which is then extracted and endocytosed for antigen processing and presentation (*Depoil et al*., 2008; *Fleire et al*., 2006; *Ketchum et al*., 2014; *McShane & Malinova*, 2022; *Spillane & Tolar*, 2017). To study the formation and organization of this specialized structure, the IS can be modeled on various antigen-bearing surfaces in vitro. While using a model APC, such as Cos-7 cells expressing specific antigens on their surface (*Fleire et al*., 2006; *Wang et al*., 2018), provides the closest equivalent to in vivo conditions, particularly in the context of germinal center responses (*Young & Brink*, 2021), they entail technical challenges for detailed analysis. As a result, many studies employ simplified setups with antigen-coated surfaces to recapitulate various aspects of the IS. Plastic or glass surfaces are the most commonly used methods, however, lipid bilayers and polyacrylamide gels can better mimic important physical properties like surface fluidity and varying stiffness, respectively (*Aceitón et al*., 2025; *McArthur et al*., 2025). Despite the critical importance of the IS for lymphocyte activation, our understanding of the underlying mechanisms and regulation remains limited. In particular, early dynamics (from first cell-cell contact to spreading) are rapid and complex, with the initial contact and receptor engagement happening within seconds and full spreading completed within minutes. Consequently, detailed study of these dynamic processes has been challenging using traditional models.

In the past, micropatterning of substrates has been used as a tool to standardize the otherwise heterogeneous shape and size of living cells, enabling more systematic and accurate analysis of biological processes, including cell geometry, membrane tension, and cytoskeletal dynamics (*Huda et al*., 2012; *Théry et al*., 2006). In immunology, micropatterns have also been used to study different aspects of T cell activation, including the effects of different co-stimulatory molecules and T cell receptor (TCR) signaling (*Motsch et al*., 2019; *Sadoun et al*., 2021; *Shen et al*., 2008). Conventional, static micropatterns are inherently limited in their ability to model highly dynamic systems. To address this limitation, we have recently developed dynamic micropatterning, which utilizes biotin–streptavidin chemistry to functionalize, on demand, the non-adhesive regions surrounding the micropatterns. This method allows precise temporal control over the triggering of cellular processes of interest, while maintaining compatibility with high-resolution microscopy for detailed spatiotemporal analysis (*Isomursu et al*., 2023).

In this work, we demonstrate the use of dynamic micropatterning to study IS formation using multiple ligands and varied pattern geometries in B and T lymphocytes. With this novel approach, we observed a surprisingly distinct localization of activated signaling proteins in the B cell synapse, with phosphorylated CD79A (pCD79A) enriched at the cell periphery, where new BCRs ligate to the antigen, while pPLCγ2 concentrated in the non-activatory area in the IS center. Notably, the use of the dynamic micropatterns enabled high resolution visualization of the earliest steps of IS formation using live imaging. This revealed an unexpected feature of B cell spreading mediated by pre existing apical, actin based veils. Altogether, our work demonstrates how dynamic micropatterns unlock new possibilities for studying early immune cell activation with both spatial precision and temporal resolution

## Results

### Dynamic micropatterns enable controlled modeling of the formation of the IS

Dynamic micropatterns were prepared by adapting the protocol from Isomursu et al., to model B cell IS formation (*Isomursu et al*., 2023) (Fig. 1A). Briefly, microscopy coverslips were coated with hydrophobic PLL-PEG-biotin, and the desired shapes were printed onto the surface via UV irradiation through a photomask, rendering the resulting micropatterned areas hydrophilic again. The micropatterns were then coated with a mixture of fibronectin and fluorescently labelled fibrinogen to promote cell adhesion. After seeding the cells onto the micropattern, a second strep-conjugated substrate can be added to functionalize the surrounding area and activate the cells. For B cells, we used Fab fragments of an anti-IgM antibody (Strep-αIgM Fab) acting as a surface-bound surrogate antigen. We and others have previously shown that αIgM Fab fragments do not activate the BCR in soluble form (*Isomursu et al*., 2023; *Volkmann et al*., 2016).

**Figure 1:**
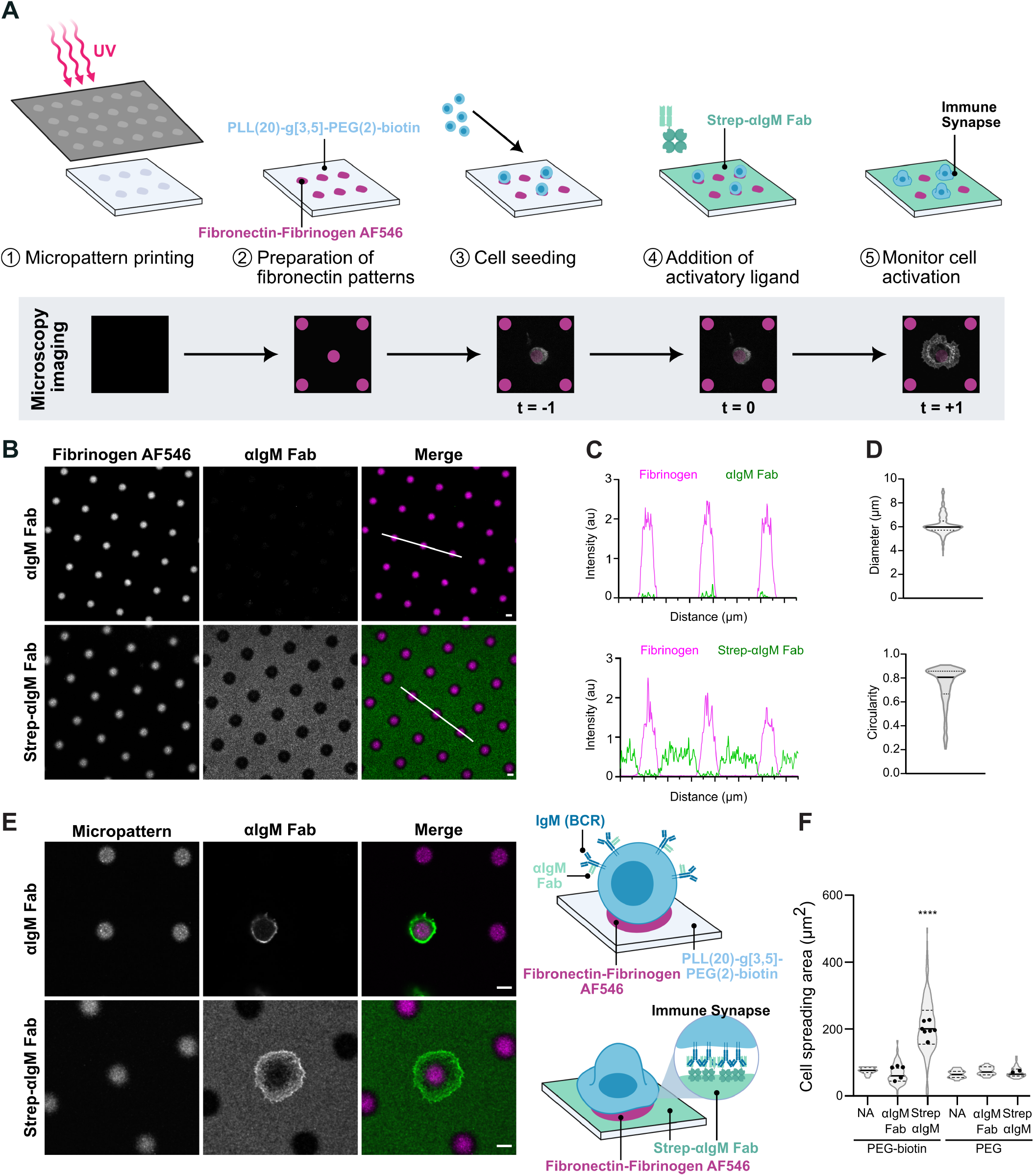
Adapting dynamic micropatterns to model the B cell IS. **A)** A schematic presentation of the production of coverslip (top) for dynamic micropatterning and representative confocal images of the B cell activation in each step of the process (bottom). **B)** Confocal representative images of 5 µm circular micropatterned coverslips with fibronectin and fibrinogen AF546 (magenta) coating and the binding of streptavidin conjugated αIgM Fab AF488. Scale bar 5 µm.**C)** The intensity of the signal from fibrinogen and αIgM Fab along the lines marked in merged images of B. **D)** Violin plot graph with mean (central line) and quartiles (dotted lines) of the diameter and circularity of the patterns as in B. **E)** Representative confocal images of A20 D1.3 B cells seeded in 5 µm circular micropatterns (magenta) for 20 minutes and in the presence of unconjugated αIgM Fab (green) (top) or streptavidin conjugated αIgM Fab (green) for 15 minutes (bottom) with schematic representation of both conditions (right). Scale bar 5 µm.**F)** The quantification of the data in (E). B cell spreading area measured by F-actin stained with phalloidin-670 during activation on PLL-PEG coated or PLL-PEG-biotin coated micropatterned coverslips. Violin plot graph with mean (central line), quartiles (dotted lines) and experiment means (dots) are shown.

First, we verified the specificity and precision of substrate binding. As expected, fluorescently labelled fibrinogen was detected exclusively on the micropatterns generated by UV irradiation through the photomask (Fig. 1B-C). Correspondingly, when streptavidin (Strep)conjugated αIgM Fab fragments were added, they bound specifically to the PLL-PEG-biotin-coated areas surrounding the fibrinogen patterns. To examine the consistency and accuracy of pattern production, we measured the diameter and circularity of the micropatterns produced through a photomask with circular dot shapes of 5 µm in diameter (Fig. 1D). The resulting patterns were largely homogeneous, with an average diameter of approximately 6 µm, indicating that the irradiated area is slightly larger than the permissive hole in the mask, a typical consequence of the thin space left between the coverslip and the photomask upon irradiation.

We went on to test the performance of dynamic micropatterns to analyze the IS formation using A20 D1.3 cells, an IgM+ murine B cell line. The cells were first allowed to adhere for 20 minutes to the circular micropatterns coated with fibronectin and fluorescently labeled fibrinogen. This substrate mixture did not induce any detectable signaling response, thereby maintaining the cells in a resting state before triggering activation (Supplementary material, Fig. S1A-B). As expected, the unconjugated αIgM Fab fragments, added to the samples, bound to the BCRs on the cell membrane but did not adhere to either the patterned or unpatterned surface (Fig. 1E, upper panel). Due to the lack of the BCR cross-linking and binding to the substrate, these unconjugated Fab fragments are unable to trigger BCR activation in solution (Fig. 1E-F). In contrast, Strep-conjugated αIgM Fab fragments robustly bound to the PLL-PEG-biotin, inducing activation as evidenced by cell spreading and immune synapse formation (Fig. 1E, lower panel and Fig. 1F). These experiments demonstrate that only the combination of a permissive PLL-PEG-biotin surface and a surface-bound activatory ligand (Strep-αIgM Fab) can trigger IS formation (Fig. 1F), highlighting the specificity and suitability of this system for modeling immune synapse formation under controlled conditions.

### Downstream BCR signaling proteins exhibit spatially distinct activation profiles in ligand-bound and unbound regions

Dynamic micropatterns offer a unique platform for directly comparing surface regions with or without BCR ligands, enabling us to investigate spatial signal segregation downstream of BCR engagement. To do this, B cells were seeded on fibronectin-coated micropatterns and activated with Strep-αIgM as described before for 0, 5, 15 or 30 minutes. Cells were fixed and stained with phospho-specific antibodies against phosphorylated CD79A (pCD79A), pPLCγ2, pBtk, and pAkt, as well as phalloidin to monitor cell spreading (Fig. 2A-C). The cells robustly spread on the Strep-αIgM ligand utilizing F-actin-based protrusions of either smooth lamellipodial or spiky filopodial phenotypes, or a mixture of these two. BCR signaling is initiated by phosphorylation of CD79A, an integral part of the BCR complex bearing the immunoreceptor tyrosine-based activation motif (ITAM) and the site of the first detectable phosphorylation in response to antigen engagement. As expected, pCD79A was enriched at the peripheral edge of the spreading cell. Among the other signaling proteins analyzed, only pBtk showed a similar enrichment pattern, co-localizing with pCD79A at the cell periphery. In contrast, pPLCγ2 consistently accumulated in the central, non-activatory areas. pAkt transitioned from initial enrichment at the cell edge into a sparse, punctate distribution all over the synaptic plane.

**Figure 2:**
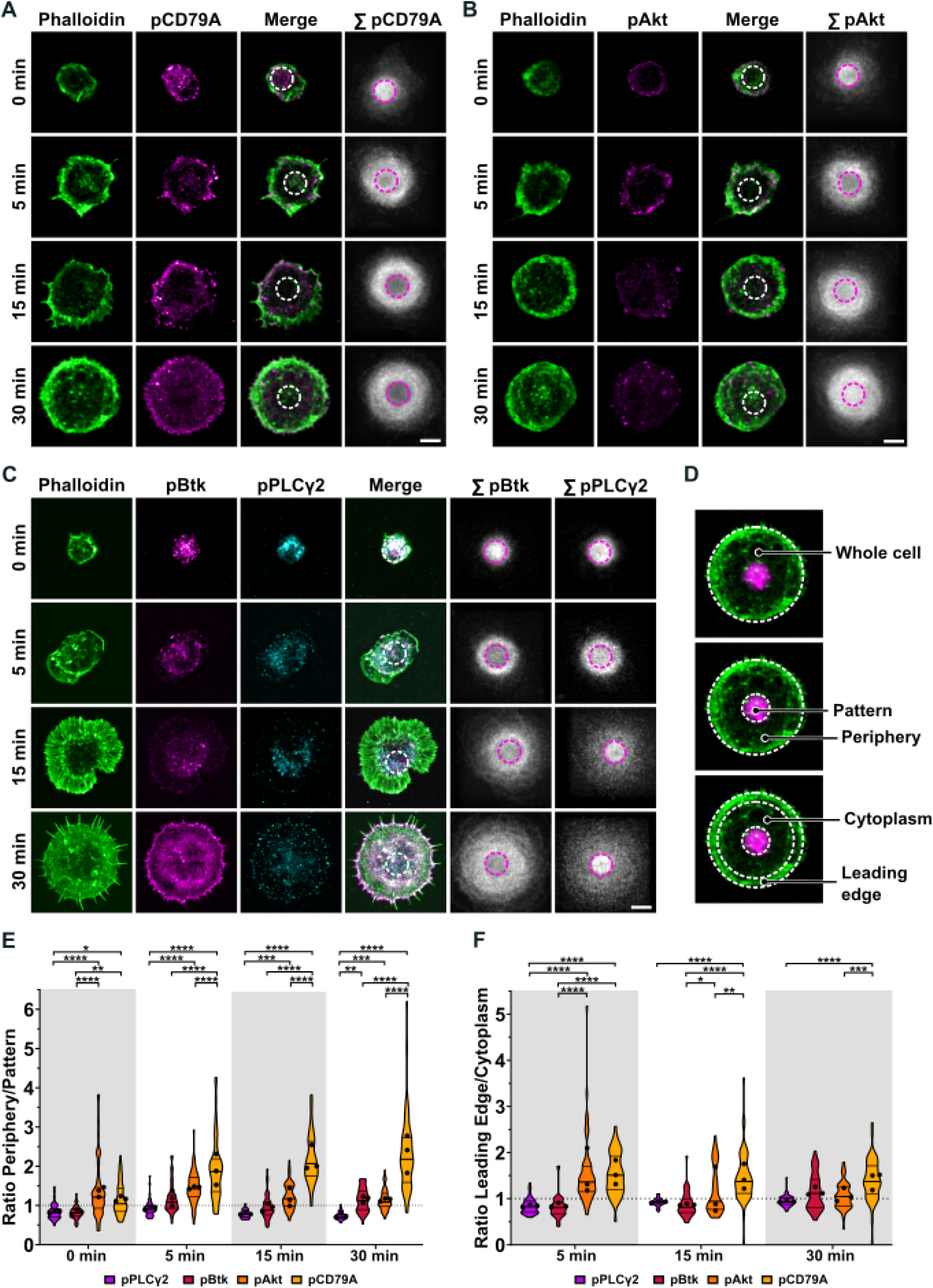
Signaling proteins downstream of the BCR show differential localization in the IS. **A-C)**) A20 D1.3 B cells were allowed to form immune synapses via first seeding them on fibronectin/fibrinogen-coated circular 5 µm (dotted white or magenta line) dynamic micropatterns for 20 minutes and then activated by the addition of Strep-αIgM Fab for 0, 5, 15 or 30 minutes. Samples were fixed and stained for immunofluorescence analysis to visualize F-actin (green) and pCD79A (magenta, A), pAkt (magenta, B), pBtk (magenta, C) and pPLCγ2 (cyan, C). Representative images of single confocal microscopy slices are shown as well as sum images of phospho-proteins (white) of all cells per condition. Scale bar 5 µm. **D)** Representation of the different regions of interest analyzed from the data in A-C (F-actin, green; fibronectin, magenta): whole cell, pattern, periphery, cytoplasm, and leading edge. **E-F)** The ratio of Periphery MFI/Pattern MFI (E) and the ratio of leading Edge MFI/Cytoplasm MFI (F) in all time points and phospho-proteins from the data in A-C. Violin plot graphs with mean (central line), quartiles (top and bottom lines) and experiment means (dots) are shown. One-way ANOVA (n=3, 15-30 cells per condition).

To validate these spatial differences, we performed signal averaging across multiple cells using the fibronectin-coated micropattern as a reference point for alignment (Fig. 2A-C, right-most panels; Supplementary material, Fig. S2). These summed projections clearly demonstrated pPLCγ2 accumulation in the central fibronectin region, whereas pCD79A was strongly excluded from it. The peripheral enrichment of pCD79A and pBtk at the leading edge was not preserved in the summed projections, due to the variability in the orientation and morphology of cell spreading. pAkt showed initial enrichment to the site of BCR-signaling but, by 30 min, the signal became uniform across the entire contact area, regardless of BCR ligand availability. While pBtk also showed exclusion from the fibronectin pattern, this effect was less pronounced than for pCD79A.

To quantitatively analyze the spatial distribution of signaling, we defined the following regions of interest: whole cell, non-activatory pattern area, cell periphery, cy-toplasm, and the leading edge (Fig. 2D). The analysis strongly supported the observations from both individual cells and signal-summed projections, supporting the differential localization of each phospho-protein during activation. Among the analyzed proteins, pCD79A showed the most distinct enrichment at the activatory area and leading edge. In contrast, pPLCγ2 was preferentially localized to the non-activatory patterned area, and pAkt and pBtk displayed intermediate behaviors (Fig. 2E-F).

Together, these data suggest a substantial spatial uncoupling of BCR downstream signaling components from the initial site of receptor engagement.

### Mechanical strain conditions signaling responses to the BCR ligation

A benefit of micropatterns is that they can be generated in various sizes and shapes, allowing for the introduction of different mechanical stresses to the cell. With this in mind, we next compared triangle-shaped patterns with 15 µm long edges to the 5 µm circular micropatterns used above (Supplementary material, Fig. S3A). Initiated from the triangles, the cells responded to the addition of Strep-αIgM with robust spreading, which, however, typically started on one of the three edges, causing more variation in cell shapes compared to the circular patterns (Fig. 3A-C; Fig. S3B). The spreading area was larger in the samples with triangles than the circles in the earliest time points, likely reflecting the larger starting point area, but already by 15 min of activation, the cells reached the same area (Supplementary material, Fig. S3C).

**Figure 3:**
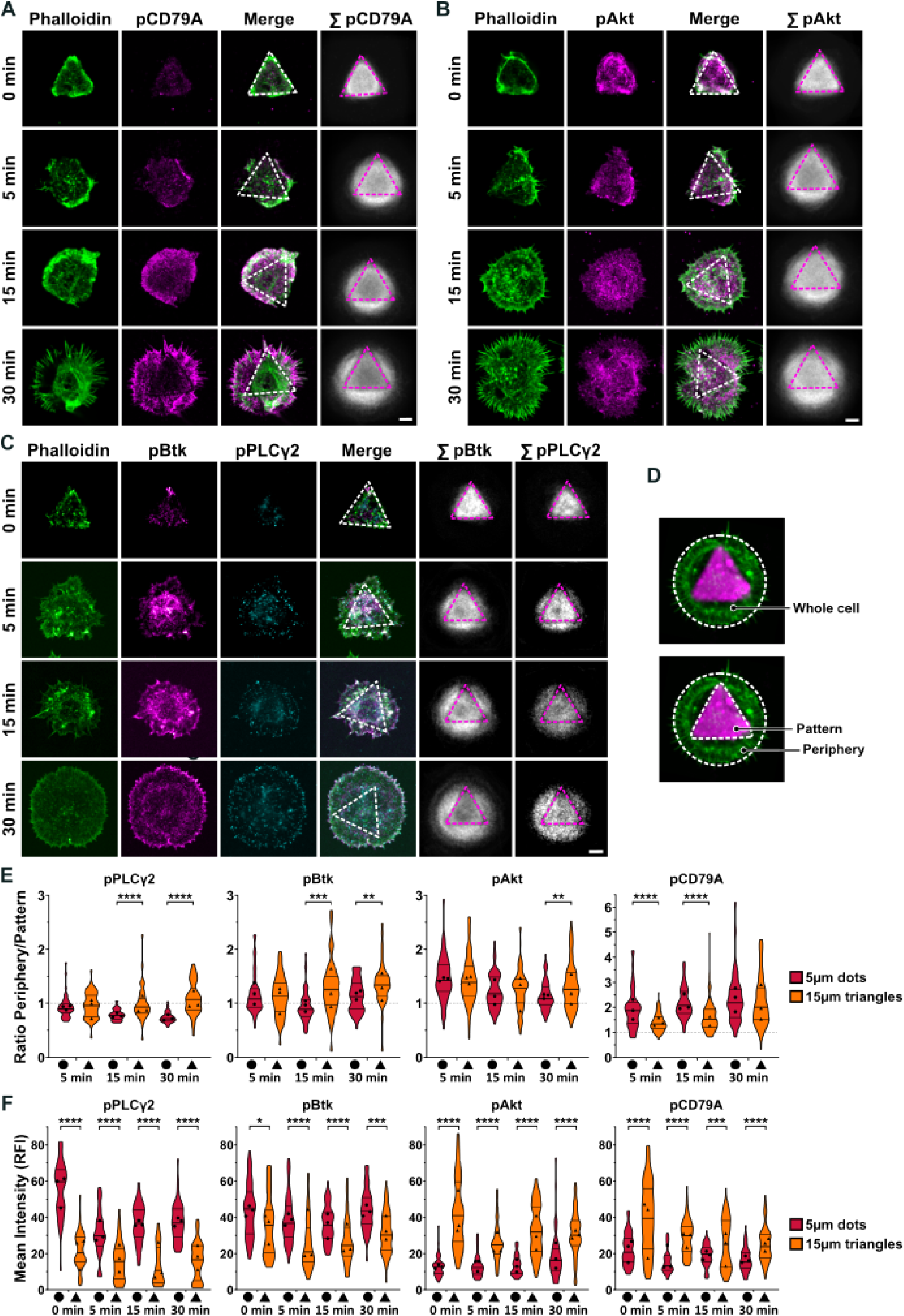
Signaling proteins downstream of the BCR show differential localization in the IS. **A-C)**) A20 D1.3 B cells were let to form immune synapses via seeding them on fibronectin/fibrinogen-coated triangular 15 µm (dotted white or magenta line) dynamic micropatterns 20 minutes and activated by the addition of Strep-αIgM Fab for 0, 5, 15 or 30 minutes. Samples were fixed and stained for immunofluorescence analysis to visualize F-actin (green) and pCD79A (magenta, A), pAkt (magenta, B), pBtk (magenta, C) and pPLCγ2 (cyan, C). Representative images of single confocal microscopy slices are shown, along with the summed images of phospho-proteins (white) from all cells per condition and staining. Scale bar 5 µm. (D) Representation of the different regions of interest analyzed in confocal images (F-actin, green) activated in 15 µm triangle micropatterns (fibronectin, magenta): whole cell, pattern and periphery. **D)** Representation of the different regions of interest analyzed in confocal images (F-actin, green) activated in 15 µm triangle micropatterns (fibronectin, magenta): whole cell, pattern and periphery. **E-F)** The ratio of periphery MFI/pattern MFI (E) and mean intensities (F) of pPLCγ2, pBtk, pAkt and pCD79A comparing the 15 µm triangle micropatterns of the data in A-C to the 5 µm circular (Figure 2 data A-C). Violin plot graphs with mean (central line), quartiles (top and bottom lines) and experiment means (dots) are shown. Unpaired t-test (n=3, 15-30 cells per condition per experiment).

To explore if the shape and size of the initial pattern could modulate the responses of the BCR signaling proteins, we seeded the cells on triangle-shaped micropatterns. In contrast to the circular patterns that the cells could fully cover without any visible strain, the triangles caused the cells to mildly stretch when adopting the shape of the non-activatory pattern. Upon activation, as seen before with the circular patterns, pCD79A and pBtk showed overall enrichment at the activatory area in the cell periphery, while pPLCγ2 and pAkt did not (Fig. 3A-C; Supplementary material, Fig. S2). However, we detected significant differences in the levels of phosphorylation in activatory versus non-activatory areas for each signaling protein when comparing the synapses initiated from triangles or circles. While pCD79A was less concentrated in the cell edges than in the synapses initiated from circles, the other phospho-proteins showed an increase in their peripheral localization. The most striking change was detected in pPLCγ2, which exhibited a clear enrichment in the non-activatory area in synapses initiated from circles, but a uniform distribution throughout different substrates in triangle-initiated synapses (Fig. 3E-F; Supplementary material, Fig. S4).

We also observed overall different signal intensities in the BCR-triggered phospho-proteins in the synapses initiated from triangles as compared to circles. When the mean intensity of the fluorescence signal was compared between the two starting shapes, two distinct groups arose; pPLCγ2 and pBtk signals were significantly reduced in cells starting from triangles, while pAkt and pCD79A signals were significantly increased (Fig. 3F). These distinct behaviors were highly significant and systematic throughout different time points. Interestingly, pAkt signal was also more homogenous throughout the activation time points, as opposed to sparse clusters detected in cells spreading from circular patterns.

Together, our analysis showed that both the distribution and abundance of phospho-proteins within the IS are influenced by the initial cell shape, as dictated by the selected micropattern geometry. This suggests that factors such as mechanical constraints and cellular symmetry may contribute to the spatio-temporal regulation of the BCR activation cascade.

### Pre-existing protrusions guide early B cell synapse formation

Dynamic micropatterns offered a unique opportunity to examine the earliest stages of IS formation, starting from a resting, non-signaling state, with high spatio-temporal resolution. The delicate F-actin-rich protrusions are difficult to fully preserve in fixed samples. This led us to use live-cell imaging, utilizing the exogenous expression of the actin reporter LifeAct-Emerald, which enables visualization of the actin cytoskeleton in real-time.

Cells were seeded onto 5 µm fibronectin/fibrinogen circles and LifeAct-Emerald fluorescence was recorded before and for 15 minutes after activation with StrepαIgM Fab. Using high-resolution 3D confocal imaging we tracked F-actin structures both at the cell’s apical side and the IS surface. We identified three distinct spreading phenotypes during IS formation: i) 47% of the cells spread with smooth lamellipodia, ii) 24% of the cells used filopodia-driven spreading, and iii) 29% of the cells generated ruffles from large, pre-existing actin protrusions from the apical side of the cells (Fig. 4A and D). Subsequently, these F-actin-based protrusions appeared to be rapidly dragged to the activatory surface to facilitate the formation of the IS (Fig. 4B). To our knowledge, such a ruffle phenotype has not been reported before—most likely because the highly dynamic, three-dimensional nature of these structures makes them difficult to visualize without the dynamic micropattern setup.

**Figure 4:**
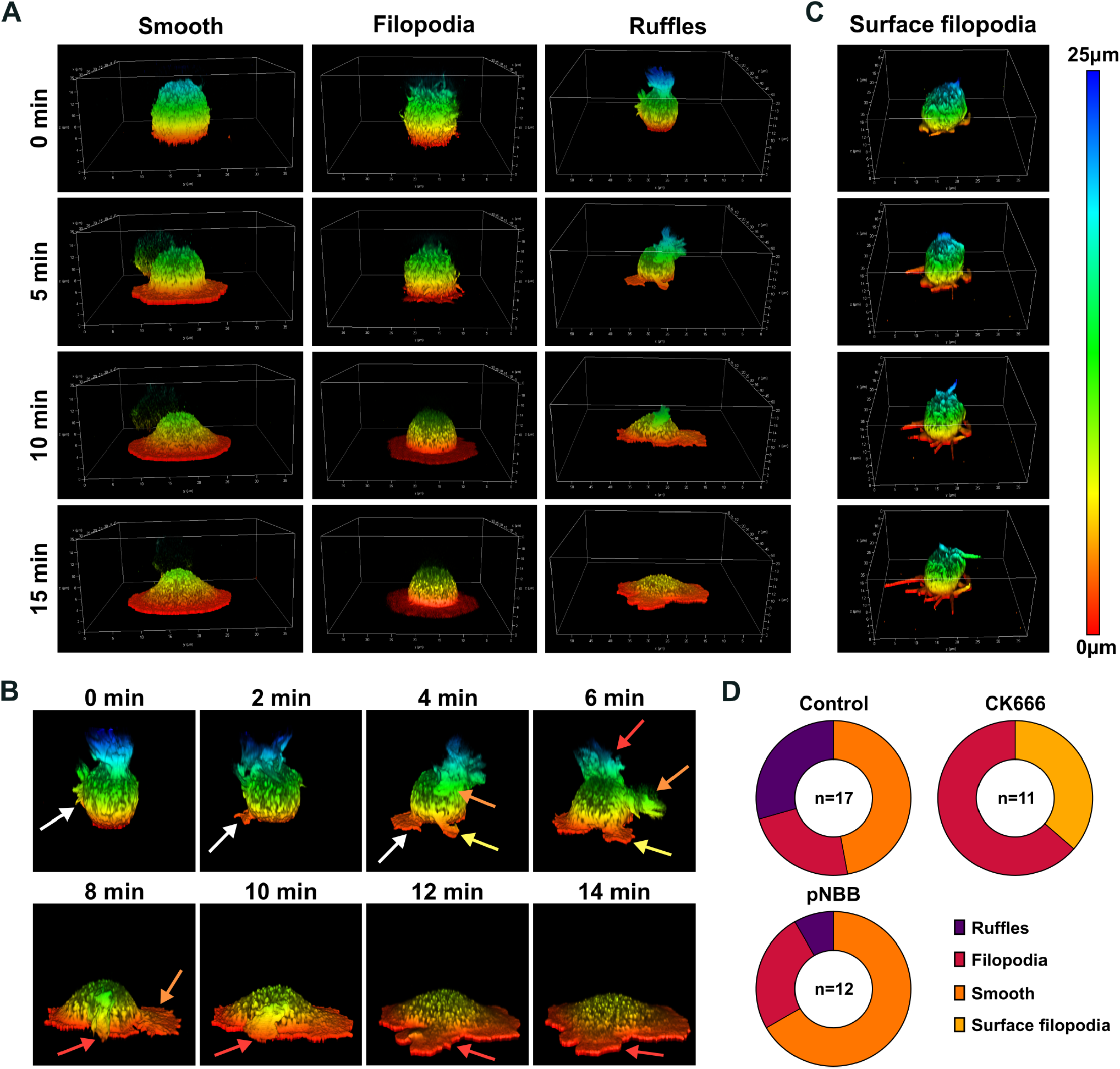
B cells utilize pre-existing F-actin protrusions to facilitate cell spreading upon IS formation. **A)** A20 D1.3 B cells, transfected with LifeAct-Emerald, were seeded on fibronectin/fibrinogen-coated circular 5 µm dynamic micropatterns for 20 minutes and activated for 15 minutes by the addition of Strep-αIgM Fab with simultaneous imaging with Leica Stellaris confocal microscope. Representative images of 3D confocal live imaging show different phenotypes: smooth spreading, filopodia-driven spreading, and actin ruffles-driven spreading. **B)** Individual frames of the representative video of a cell with the actin ruffle phenotype seen on panel A (colored arrows point to individual ruffles). **C-D)** The cells were let to form IS as in A, with the addition of the actin cytoskeleton inhibitors CK666 to inhibit Arp2/3 and para-nitro-Blebbistatin to inhibit myosin II. A representative cell, treated with CK666, showing the surface filopodia-driven spreading (C). **D)** Pie charts of the n-Dumber of cells showing each phenotype in all conditions. n=11-17 cells per condition.

To understand the mechanisms behind these three cellular behaviors, we treated the cells with inhibitors of key cytoskeletal components: CK666 (an Arp2/3 inhibitor), which blocks branched actin formation required for lamellipodia, and para-nitro-blebbistatin (a Myosin II inhibitor), which impairs actomyosin contractility. When Arp2/3 was inhibited, actin ruffles and smooth, lamellipodial spreading disappeared, leaving the cells to rely only on filopodia to spread and engage with the antigen. The inhibitor treated cells showed two distinct subphenotypes: 64% of the cells showed filopodia similar to the non-treated cells, and 36% showed so-called surface filopodia that extended only along the surface-attached side of the cells (Fig. 4C), while filopodia in the control cells grow from any area of the cell and then move to contact the activatory surface and fuse with the leading edge. Conversely, when Myosin II was inhibited, the most prevalent phenotype was the smooth spreading, and also the number of ruffling cells decreased considerably (Fig. 4D).

The changes in the prevalence of different phenotypes demonstrated the differential involvement of these cytoskeletal components in IS formation. Specifically, the cell spreading utilizing pre-existing actin ruffles depended on both Arp2/3 activity, likely to create the ruffles, and on myosin-driven contractility, probably to pull the ruffles toward the IS.

### Dynamic micropatterns support modeling of the immune synapse across cell types and ligands

Finally, to explore the wider applicability of our approach, we expanded the use of dynamic micropatterns to model more complex activation scenarios, including multiligand B cell stimulation and T cell synapse formation.

When B cells interact with an APC to form an IS, in addition to BCR-antigen binding, the integrin LFA-1 on the B cells binds to ICAM-1 on the APC surface (*Carrasco et al*., 2004; *Wang et al*., 2022). We modeled this interaction by adding ICAM-1 as a second Strep-conjugated ligand into our dynamic micropatterns. Activation was triggered by adding Strep-αIgM or StrepICAM-1 alone, or both at ratios of 2:1, 1:1, and 1:2. Cell spreading was analyzed alongside pCD79A levels (Supplementary material, Fig. S5A). Stimulation with ICAM-1 alone did not induce B cell spreading. Compared to Strep-αIgM alone, the 1:2 ratio of Strep-αIgM to Strep-ICAM-1 resulted in increased cell spreading, although with reduced overall pCD79A levels (Supplementary material, Fig. S5B-C). Conversely, the peripheral localization of pCD79a was enhanced by lower amounts of Strep-ICAM-1 (Supplementary material, Fig. S5D).

To model the T cell immune synapse, Jurkat T cells were seeded on fibronectin/fibrinogen-coated micropatterns and activated by the addition of Strep-αCD3 and Strep-αCD28 for 0 and 30 minutes. To accommodate the larger size of Jurkat cells, we used 10 µm circular micropatterns. We assessed cell spreading and phosphorylation of Akt, a signaling protein shared across B and T cell activation cascades. Upon activation, T cells spread to form synapses, similarly to B cells (Supplementary material, Fig. S5E-F). As with B cells, pAkt mainly localized to the cell periphery; however, in T cells, the signal was more evenly distributed and lacked the punctate appearance seen in B cells (Supplementary material, Fig. S5E, G). The modest level of spreading we detected for Jurkat cells could be due to the stimulatory signals the cells receive also on their apical side, in the form of TCR cross-linking Strep-αCD3 in solution. We propose that using monovalent, non-stimulatory αCD3 Fab fragments (*Minguet et al*., 2007) as Strep-linked stimulatory surrogate antigen, analogous to B cells, would enhance the response of Jurkats to form the IS in this system.

Together, these results highlight the versatility of dynamic micropatterns for modeling immune synapse formation across cell types and ligand combinations.

## Discussion

Here, we adapt the recently developed dynamic micropatterning approach (*Isomursu et al*., 2023) to study IS formation with high spatial and temporal resolution. This method enables the simultaneous analysis of IS regions with or without BCR engagement within a single cell. The method is also unique in allowing high-resolution imaging of the earliest steps of lymphocyte activation on antigen-presenting surfaces. Our results reveal that activated signaling proteins exhibit distinct spatial distributions relative to activatory or non-activatory regions of the IS. Live-cell imaging further uncovered that, in addition to de novo actin polymerization, B cells repurpose preexisting actin-rich protrusions as building material to facilitate energy-efficient spreading during IS formation.

When B cells form an IS on the surface of APCs, the recognized antigen is presented in a soft and laterally mobile membrane environment. Detailed study of IS on the native APC surface is technically challenging, particularly with microscopy-based approaches, and difficult to control. Different in vitro model systems have been developed to mimic IS formation, each with its own advantages and disadvantages, often resulting in a trade-off between physiological relevance and analytical power. For example, planar lipid bilayer systems have been developed to provide membrane fluidity (*Dustin*, 2009)), while soft substrates simulate the mechanical properties of APCs (*Aceitón et al*., 2025; *Zeng et al*., 2015). By contrast, glass is highly rigid and non-fluid, but highly amenable to highresolution imaging and diverse functionalization strategies. Using dynamic micropatterns, we achieve a high level of spatial and temporal control over ligand presentation, and the possibility to model the effects of different ligands in a straightforward manner, while remaining compatible with diverse imaging modalities, including super-resolution microscopy. This approach also enables systematic exploration of co-stimulatory and co-inhibitory signals at the IS, as demonstrated with the addition of ICAM-1 in our experiments. The main limitation of the system is its relatively low throughput, as not every pat-tern binds a cell, resulting in modest cell numbers per coverslip.

In addition, different cell lines were found to present different compatibilities with the system. For example, Raji human B cell line showed poor adhesion to the micropatterns, potentially due to interference from the surrounding PLL-PEG matrix. When we analyzed the localization of phosphorylated signaling molecules (pPLCγ2, pBtk, pAkt, and pCD79A) within the IS, we expected these signals to co-accumulate at the site of active BCR engagement. Instead, we observed distinct and non-overlapping patterns (Fig. 2A-C). pCD79A was enriched at the leading edge, consistent with this region being the site of new BCR-antigen engagement as the cell spreads to encounter new antigens. The decrease in signal toward the cell body likely reflects phosphatase-mediated negative feedback(*Doody et al*., 1995; *Gold et al*., 1991; *Wienands et al*., 1996). We also found pBtk mainly in activatory regions, highly enriched at the leading edge, aligning with its known role in actin remodeling via WASP(*Bhanja et al*., 2022; *Sharma et al*., 2009). Interestingly, pAkt showed only a slight preference for activatory areas, while pPLCγ2 strongly favored the non-activatory area of the IS when cells were seeded on the fibronectin/fibrinogen-coated circular patterns. Notably, on larger, triangular patterns, pPLCγ2 appeared more evenly distributed.

These results indicate that downstream BCR signaling rapidly diverges spatially within the IS, despite their tight connections such as shared binding partners or common complexes reported to these proteins. For example, both PLCγ2 and Btk have been found to form a complex with BLNK downstream of CD79A activation (*Fu et al*., 1998; *Hashimoto et al*., 1999; *Tanaka & Baba*, 2020), yet they show opposite localization preferences regarding the BCR ligation status. Using fluid planar lipid bilayers as the antigen-presenting substrate, B cells as well as T cells, have been shown to form “signaling microclusters”, where both receptor-antigen complexes and the proximal signaling molecules co-accumulate. However, the physiological plasma membrane of the APC presents variable diffusion for different proteins (*Comrie et al*., 2015), and microclusters become less and less prominent as substrate fluidity decreases (*Ketchum et al*., 2014; *McArthur et al*., 2025).

Substrate mechanics introduce an additional layer of regulation. Mechanosensing of the activatory surface stiffness has emerged an important factor in B cell activation, influencing processes such as IS formation (*Aceitón et al*., 2025), polarization (*Pinon et al*., 2022), activation threshold (*Brooks et al*., 2023) or antigen extraction (*Spillane & Tolar*, 2017). Here, we used small circular and larger triangular micropatterns to simulate different membrane tensions at the onset of activation. When comparing the sig-naling protein distributions between these two conditions, we observed shifts in both the spatial preference and overall intensity of signaling proteins. PLCγ2 and Btk activation were reduced on triangular patterns, whereas Akt and CD79A activation increased. This pattern indicates that various branches of BCR signaling respond differently to changes in membrane tension. For instance, it is known that mechanical stretching can induce phosphorylation of Akt in a PI3K-independent manner in other cell types (*Kippenberger et al*., 2005). Conversely, PLCγ2 and Btk have been linked to the ability of B cells to distinguish between soft and stiff substrates (*Shaheen et al*., 2017). Although the mechanisms of mechanosensing and its effects in B cells are still poorly understood, it could be speculated that the effects we observe are related, for example, to calcium signaling, as mechanosensing ion channels can influence calcium levels in B cells when in contact with glass surfaces (*Kwak et al*., 2023).

A key advantage of dynamic micropatterns is the precise synchronization of activation of cells pre-bound to non-activatory patterns, allowing high-resolution live imaging of the earliest stages of IS formation. Live imaging also enables visualization of delicate actin-based protrusions that are often lost during fixation. We effectively observed IS formation in 3D to examine the dynamics of the actin cytoskeleton during early activation. We identified three different phenotypes, characterized by the predominant actin structures formed during activation: smooth spreading, filopodia-driven spreading, and asymmetric spreading utilizing pre-existing apical actin ruffles. In the ruffle-based mode, cells appeared to recycle pre-existing F-actin and cell membrane as a whole to facilitate synaptic spreading in an energy-efficient manner. To our knowledge, this phenotype—dependent on Arp2/3 and myosin II contractility—has not been previously described. Its discovery underscores the unique ability of dynamic micropatterns to capture the earliest, mechanically and spatially complex phases of IS formation. Overall, these findings position dynamic micropatterns as a powerful platform to dissect the spatial, mechanical, and temporal mechanisms of immune synapse formation with a level of precision that surpasses existing model systems.

## Materials and Methods

Detailed information of antibodies, reagents and consumables used can be found in Supplementary material, Tables S1 and S2.

### Cell lines

A20 D1.3 mouse lymphoma cells (*Williams et al*., 1994), which express a hen egg lysozyme specific IgM BCR, were kind gifts from Prof Facundo Batista (the Ragon Institute of MGH, MIT and Harvard, USA). Cells were cultured in complete RPMI (cRPMI) (RPMI 1640 with 2.05 mM L-glutamine supplemented with 10% fe-tal calf serum (FCS) 5 µM 1468 B-mercaptoethanol, 4 mM L-glutamine, 10 mM HEPES and 100 U/ml penicilin/streptomycin) and maintained at +37 °C and 5% CO2. Jurkat human T cells were a kind gift from Maija Hollmén (Turku Bioscience, University of Turku and Åbo Akademi University, Finland). Cells were cultured in complete RPMI (RPMI 1640 with 2.05 mM L-glutamine supplemented with 10%fetal calf serum (FCS, 10 mM HEPES and 100 U/ml penicilin/streptomycin, 1 mM sodium pyruvate, 1.5 g/L sodium bicarbonate) and maintained at +37 °C and 5% CO2.

### Western blotting

48-well plates were coated with 200 µl of 10 µg/ml of goat anti-mouse IgM F(ab)2 fragments (Jackson ImmunoResearch, 115-006-020), 10 µg/ml fibronectin (F4759, Sigma Aldritch) or 10 µg/ml combination of fibronectin and fibrinogen AF546 (F13192, Thermo Fisher Scientific) for 2 hours at 37 °C. A20 D1.3 B cells were starved for 15 min in RPMI medium supplemented with 0.5%FCS at 3*106 cells/ml, 500.000 cells were seeded per well and stimulated for 15 or 30 min at +37 °C and 5% CO2. After stimulation, cells were instantly lysed with 5× SDS lysis buffer (to a final concentration of 62.5 mM Tris–HCl, pH 6.8, 2% SDS, 10% glycerol, 100 mM b-mercaptoethanol, and bromophenol blue) and sonicated for 5min (high power, 30 s on/off cycle, Bioruptor plus, Diagenode). Lysates were run on 10% polyacrylamide gels and transferred to PVDF membranes (Trans-Blot Turbo Transfer System, Bio-Rad). Membranes were blocked with 5% BSA in TBS (TBS, pH7.4) for 1 h and incubated with 1:1000 anti-phosphotyrosine (05-321, Merck) in 5% BSA in TBST (TBS, 0.05% Tween-20) at +4 ºC, overnight. After washing, horseradish peroxidase (HRP) conjugated antibodies (1:10,000) were incubated for 1 hour at RT in 5% milk in TBST. After final wash, membranes were visualized with Immobilon Western Chemiluminescent HRP Substrate (WBKLS0500, Millipore) in a ChemiDoc MP Imaging System (Bio-Rad). Blots were stripped with 25 mM glycine-HCl buffer, 2% SDS, pH 2.5, for 10 min at +37 ºC, blocked, and re-probed with anti-β-actin antibody (HRP) (1:20,000; AC-15, Abcam, ab49900). Washing steps were done in 10 ml of TBST for 5 × 5 min. For quantification, Images were background subtracted, and the raw integrated densities of the lanes (pTyr) were normalized to corresponding total β-actin densities in ImageJ.

### Ligand conjugation

Anti-IgM Fab (Jackson ImmunoResearch, 115-007-020), ICAM-1 (BioLegend, 553004), anti-CD3 (Invitrogen, 16-0037-85) and anti-CD28 (Invitrogen, 16-0289-81) were conjugated to streptavidin using the Streptavidin Conjugation Kit Lightning-Link (Abcam, ab102921) following manufacturer instructions.

### Micropatterned coverslip production

13 mm or 25 mm diameter 0.16-0.19 mm thickness round coverslips (Epredia) were prepared as previously described (*Iso-mursu et al*., 2023). Coverslips are first cleaned with 70% and 94% ethanol consecutively and then in a UV oven (UVO-cleaner 342-220, Jelight Company; low-pressure mercury vapor lamp, fused quartz, λ = 185 and 254 nm, 3033 mW cm2 at 254 nm with the distance of¼) for 15 minutes when all the ethanol has evaporated from the glass surface. Coverslips were coated in a wet bed with 0.1 mg/ml PLL(20)-g[3.5]-PEG(2)/PEG(3.4)-biotin(50%) (SuSoS, Switzerland) in dH2O for one hour at room temperature, washed twice in PBS and once in dH2O. Coated coverslips were then patterned by using quartz/chromium photomasks with areas for 5 µm and 10 µm diameter circular micropatterns and 15 µm side equilateral triangular micropattern (Delta Mask, Netherlands). Photomasks were cleaned with dH2O, 70% and 94% ethanol, dried using airflow and then cleaned in the UV oven for 5 minutes. Coverslips were placed on the photomask, inverted on a drop on dH2O and exposed to UV light for 6 minutes. The patterned coverslips were stored at +4 °C until use.

A maximum of 24 hours before use of patterned coverslips were coated with 10 µg/ml fibronectin (Sigma-Aldritch, F4759) and and 1:200 fibrinogen-AlexaFluor 546 (Thermo FIsher Scientific, F13192) in PBS for 1 hour at room temperature, washed twice in PBS and stored in PBS at +4 °C until use.

### IF sample preparation

Micropatterned 13 mm coverslips were prepared as previously described and placed in 24-well plates with 300 µl of starvation RPMI (sRPMI) (RPMI 1640 with 2.05mM L-glutamine supplemented with 2% fetal calf serum (FCS) 5 µM 1468 B-mercaptoethanol, 4 mM L-glutamine, 10 mM HEPES and 100 u/ml penicillin/streptomycin). A20 D1.3 cells were centrifuged at 700 rpm for 5 minutes and resuspended at 106 cells/ml in sRPMI. 200.000 cells were seeded per coverslip for 15 minutes at +37 °C and 5% CO2. After seeding coverslips are washed twice with sRPMI to remove unbound cells. Cells seeded in 5 µm dot micropatterns were directly activated using 0.5 µl of Strep-anti-IgM Fab (stock 1 mg/ml), 1 µl of Strep-ICAM (stock 0.5 mg/ml) or a mix of both at different ratios (2:1, 1:1 and 1:2) per coverslip and cells seeded in 15 µm triangle micropatterns are allowed to get in the shape of the pattern for one hour at +37 °C and 5% CO2 and then activated. After 5, 15 or 30 minutes of activation cells were fixed using 4% PFA 10 minutes RT and washed twice in PBS.

For Jurkat cells preparation of micropatterned coverslips and seeding of cells in 10 µm dot micropatterns followed the same procedure but was done using complete RPMI. Cells were activated using 0.5 µl of Strep-anti-CD3 (stock 1 mg/ml) and 0.5 µl of Strep-anti-CD28 (stock 1 mg/ml). After 30 minutes of activation cells were fixed using 4% PFA 10 minutes RT and washed twice in PBS.

In a wet chamber, samples were permeabilized and blocked (5% donkey serum, 0.3% TritonX-100 in PBS) for 20 minutes RT. After blocking cells were incubated with primary antibodies and ActiStain-488 Phalloidin (Cytoskeleton, PHDG1-A) or Phalloidin-iFluor-405 (Abcam, ab176752) at 1:200 in staining buffer (1% BSA, 0.3% TritonX-100 in PBS) at +4 °C overnight, washed in PBS to remove unbound antibodies and the incubated in Donkey anti-Rabbit IgG AF647 (Invitrogen, A31573) secondary antibody at 1:500 in PBS 45 minutes RT, washed in PBS and the mounted using Fluoromount G with (Invitrogen, 00495952) or without DAPI (Southern Biotech, 0100-01).

### Pattern quality control/Characterization

5 µm dot micropatterned 13 mm coverslips were produced as previously described, and anti-IgM Fab (Jackson ImmunoResearch, 115-007-020) or anti-IgM Fab conjugated with Streptavidin was added, washed twice in PBS and stained with Donkey anti-Goat IgG AF488 (Thermo Fisher Scientific, A-11055) 1:500 in PBS for 1 hour RT in a wet chamber.

For the control of B cell activation in different surfaces A20 D1.3 cells were seeded as previously described in coverslips coated with PLL-PEG-biotin (SuSoS, Switzerland) or PLL-PEG and micropatterned as previously described. Cells were then activated with anti-IgM Fab (Jackson ImmunoResearch, 115-007-020) or Strep conjugated anti-IgM Fab for 15 minutes at +37°C and 5% CO2, cells were then fixed with 4% PFA 10 minutes RT, washed twice in PBS, blocked (5% donkey serum, 0.% TritonX-100 in PBS) for 20 minutes RT, stained with Donkey anti-Goat IgG AF488 (Thermo Fisher Scientific, A-11055) 1:500 in PBS for 1 hour RT in a wet chamber, washed twice with PBS and mounted using Fluoromount G (Southern Biotech, 0100-01).

### Live imaging

A20 D1.3 cells were transfected as previously described (*Šuštar et al*., 2018). Cells suspended in 2S transfection buffer (5 mM KCl, 15 mM MgCl2, 15 mM HEPES, 50 mM sodium succinate, 180 mM Na2HPO4/NaH2PO4 pH 7.2) are transfected with 1 µg per 106 cells of plasmid and electroporated using an AMAXA electroporation machine (program X-005, Biosystem) in 0.2 cm gap electroporation cuvettes. LifeAct-Emerald plasmid was a kind gift from Guillaume Jaquemet (Åbo Akademi University, Turku, Finland).

12 to 24 hours after transfection micropatterned 25 mm coverslips in cell chambers (Invitrogen, A7816) are seeded as previously described with transfected cells. Just before imaging unbound cells are washed with chamber buffer (0.5% FCS, 2 mM Mg2+, 0.5 mM Ca2+, 1 g/L Dglucose in PBS). Imaging is started in Leica Stellaris 8 confocal microscope (3D imaging) and after one minute activation is added as previously described without stopping the imaging and activation is recorded for 15 minutes. When used, cytoskeleton inhibitors are added for pretreatment before imaging (CK666 100 µM 30 minutes; para-nitro-Blebbistatin 25 µM 30 minutes).

### Image acquisition

*SDCM fixed samples*. Images were acquired with a 3i CSU-W1 spinning disk equipped with 405, 488, 561 and 640 nm solid state lasers and 445/45, 525/30, 617/37 and 692/40 nm filters and 63x Zeiss Plan-Apochromat objective. Photometrics Prime BSI sCMOS (2048 × 2084 pixels, 1 × 1 binning) camera was used.

*Leica Stellaris live imaging*. Images were acquired with a Leica Stellaris 8 Falcon FLIM equipped with a white light laser and a 405 diode laser and a HC PL APO 63x/1,40 OIL CS2 objective. Power HyD S SP pos 2, Power HyD S SP Core Unit Pos 3, Power HyD X SP pos 4 and Power HyD R SP pos 5 detectors were used.

### Image analysis and processing*SDCM*

IS plane images and polarization assay images were analyzed using Fiji ImageJ. IS plane images were thresholded using the automatic threshold Huang 2 in the F-actin and the fibronectin-fibrinogen channels to create the necessary regions of interest (ROI) for the analysis of the mean fluorescent intensity (MFI) of the different phospho-proteins in the full cell, periphery, pattern, leading edge and cytoplasm as well as to calculate the ratios between regions. Leading edge to cytoplasm analysis was not performed on triangle patterns as a large fraction of the cells did not form such well-organized lamellipodia as the cells spreading from the circles, inhibiting confident thresholding of the leading edge for automated analysis. Sum images were created by aligning all images per condition by using Fast4Dreg alignment on the micropattern channel, applying drift correction to all channels, normalizing signal and then producing a sum image of the stack.

*Leica Stellaris*. Live imaging was processed using Las X adaptive Lightning processing adjusted for water medium to enhance signal. Phenotypes were described and quantified manually by the observation of the most prominent actin features during activation and spreading.

### Statistical analysis and illustrations

Statistical significance was calculated using ordinary one-way ANOVAs or unpaired Student’s t-test assuming normal distribution of the data. Statistical significances are denoted as follows: *P < 0.05, **P < 0.01, ***P < 0.001, ****P < 0.0001. Statistical analysis and graphs were generated in GraphPad Prism 8 and illustrations using BioRender or Inkscape v1.2. Figure formatting was also performed using Inkscape v1.2.

## Author Contributions

B.T-G., S.H-P., and P.K.M designed and conceptualized the research; B.T-G., S.H-P. and E.K. optimized the method and performed quality control; B.T-G. performed research, analyzed data, and prepared figures; J.I. contributed new tools; P.K.M. and J.I. provided resources and obtained funding;and B.T-G. and P.K.M. wrote the paper with input and review from all authors.

## Acknowledgments

We are thankful to Diogo Cunha, Laura Grönfors and Thomy Garnier for technical assistance, Dr Aleksi Isomursu for technical advice, and Dr Guillaume Jacquemet for the advice on the data analysis. Turku Bioscience Cell Imaging and Cytometry Core and Biocenter Finland are acknowledged for providing research infrastructures. This work was supported by the Research Council of Finland’s Flagship InFLAMES (decision numbers: 337530 and 357910) and project funding (decision numbers: 296684, 327378, and 339810 to P.K.M.), Sigrid Jusélius foundation (to P.K.M.), Magnus Ehrnrooth foundation (to B.T-G.), University of Turku Graduate School (to B.T-G.), Turun Yliopistosäätiö (to S.H-P.) and the Finnish Cultural Foundation (to S.H-P).

## Supplementary Information

- Supplementary Figures 1-5
- Supplementary Tables 1-2

**Figure S1:**
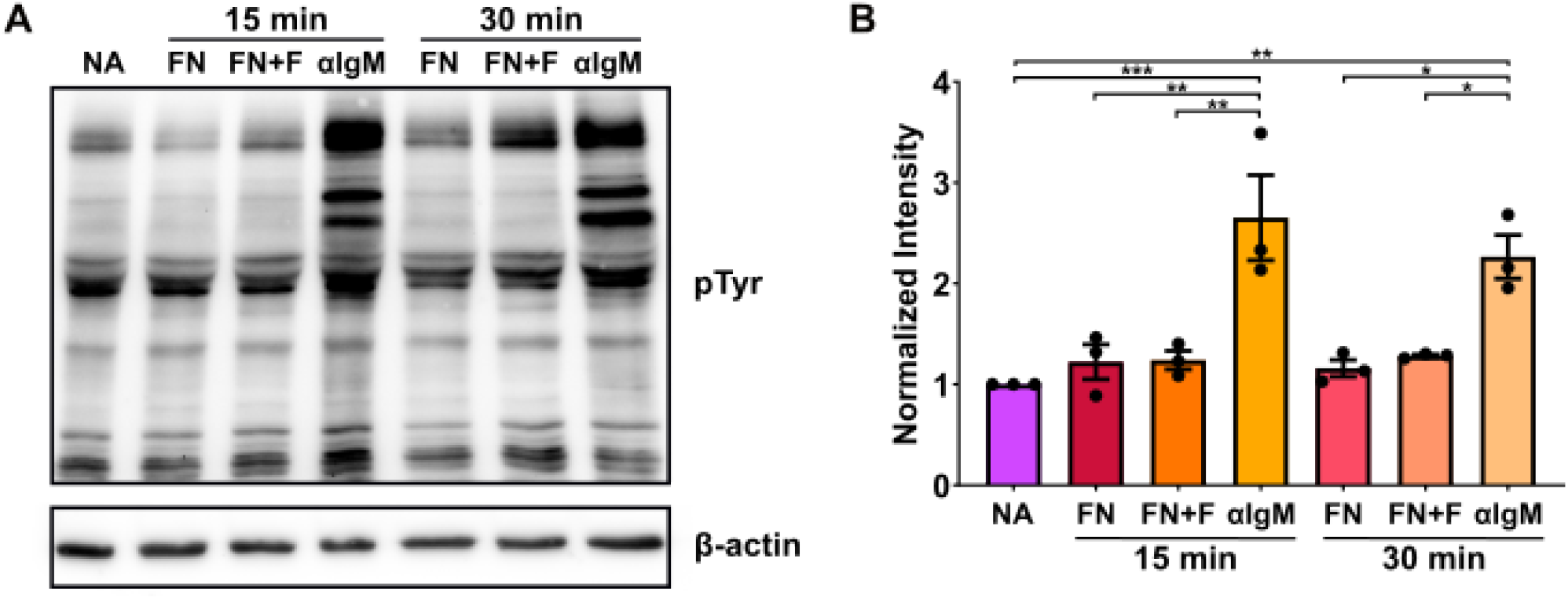
B cells are not activated by the combination of fibronectin and fibrinogen. (**A**) B cells were seeded on glass coverslips left uncoated (NA) or coated with fibronectin (FN), fibronectin and fibrinogen AF546 (FN+F) as used for dynamic micropatterns, or αIgM Fab2 (αIgM). After 15 or 30 minutes, the samples were lysed and immunoblotted with HRP-anti phospho-tyrosine and anti-β-actin as loading control. (**B**) Bar graph of the quantification of normalized intensity with mean, SEM and individual experiment values (dots). Values are shown as change compared to non-activated cells after normalization of values using the β-actin loading control. One-way ANOVA (n=3 experiments).

**Figure S2:**
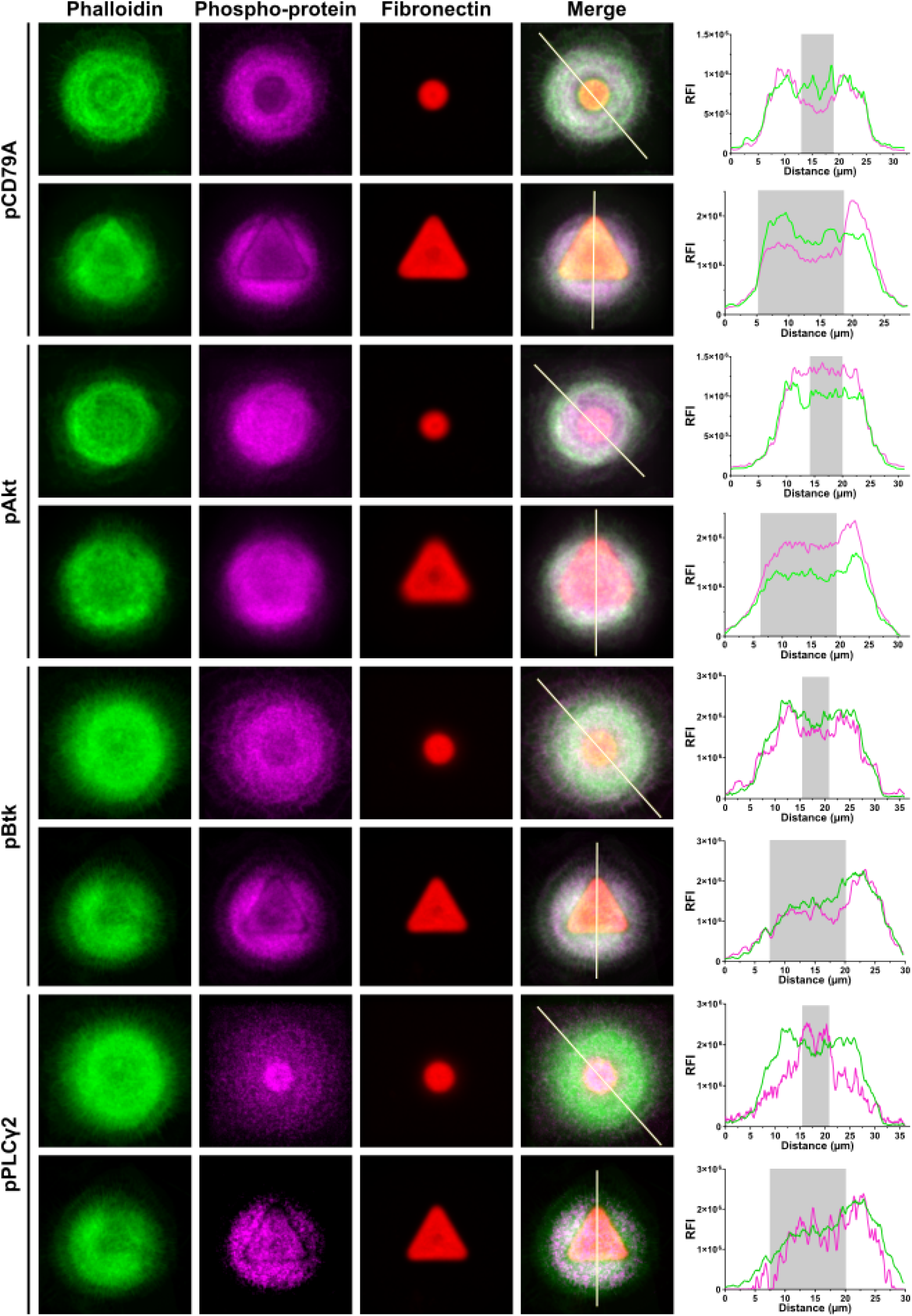
Analysis of different localization patterns on dynamic micropatterns. All images of single confocal microscopy slices of B cells activated with Strep-αIgM Fab for 30 minutes on 5 µm circular or 15 µm triangle dynamic micropatterns, fixed and stained for F-actin (green) and phospho-proteins (magenta), were aligned using the fibronectin/fibrinogen-AF546 signal. Summed images of all cells per condition were created using Fiji. Histograms represent RFI of F-actin (green) and phospho-proteins (magenta) along the white line drawn on merge images, grey background area marks the section that transverses the micropattern.

**Figure S3:**
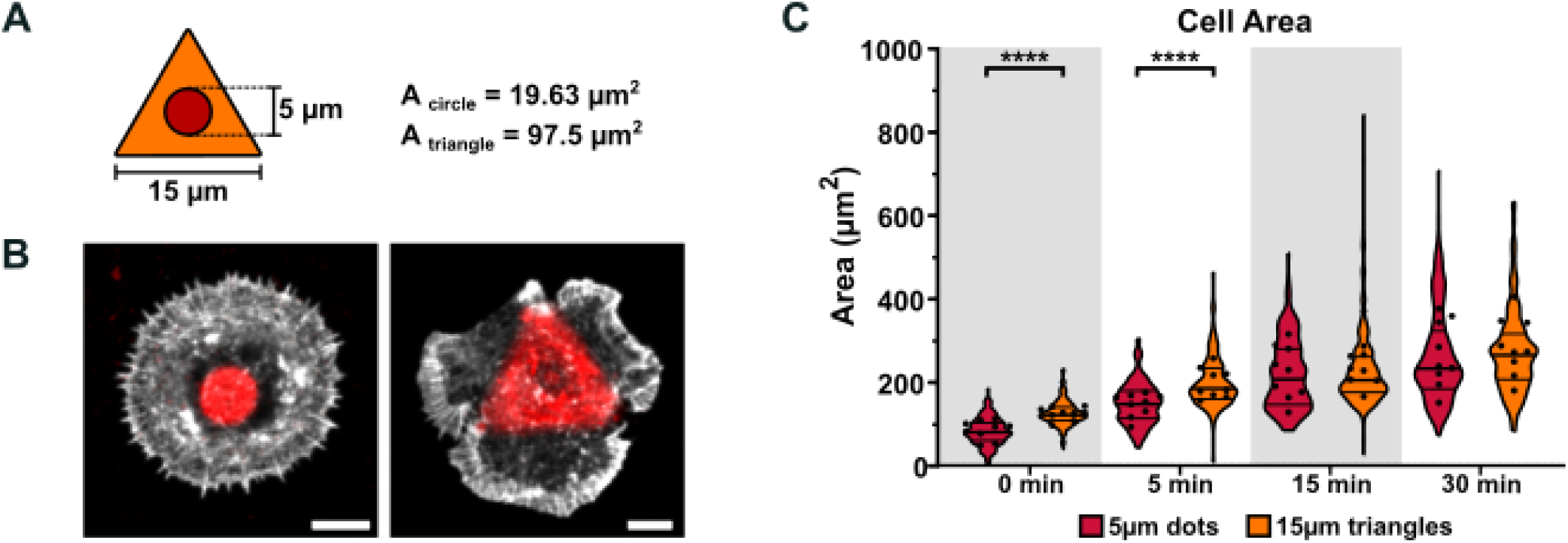
Triangular micropatterns pre-shape B cells compared to circular micropatterns. (**A**) Representative drawings of the sizes of 5 µm circular and 15 µm triangle micropatterns to scale. (**B**) Representative confocal images of A20D1.3 mice B cells let to activate for 30 min from 5 µm circular or 15 µm triangle micropatterns (fibronectin, red) fixed and stained for F-actin (white). Scale bar 5 µm. (**C**)Violin plot graphs with mean (central line), quartiles (top and bottom lines) and experiment means (dots) of cell area in the immune synapse measured by F-actin staining. One-way ANOVA (n=9, 15-30 cells per condition per experiment).

**Figure S4:**
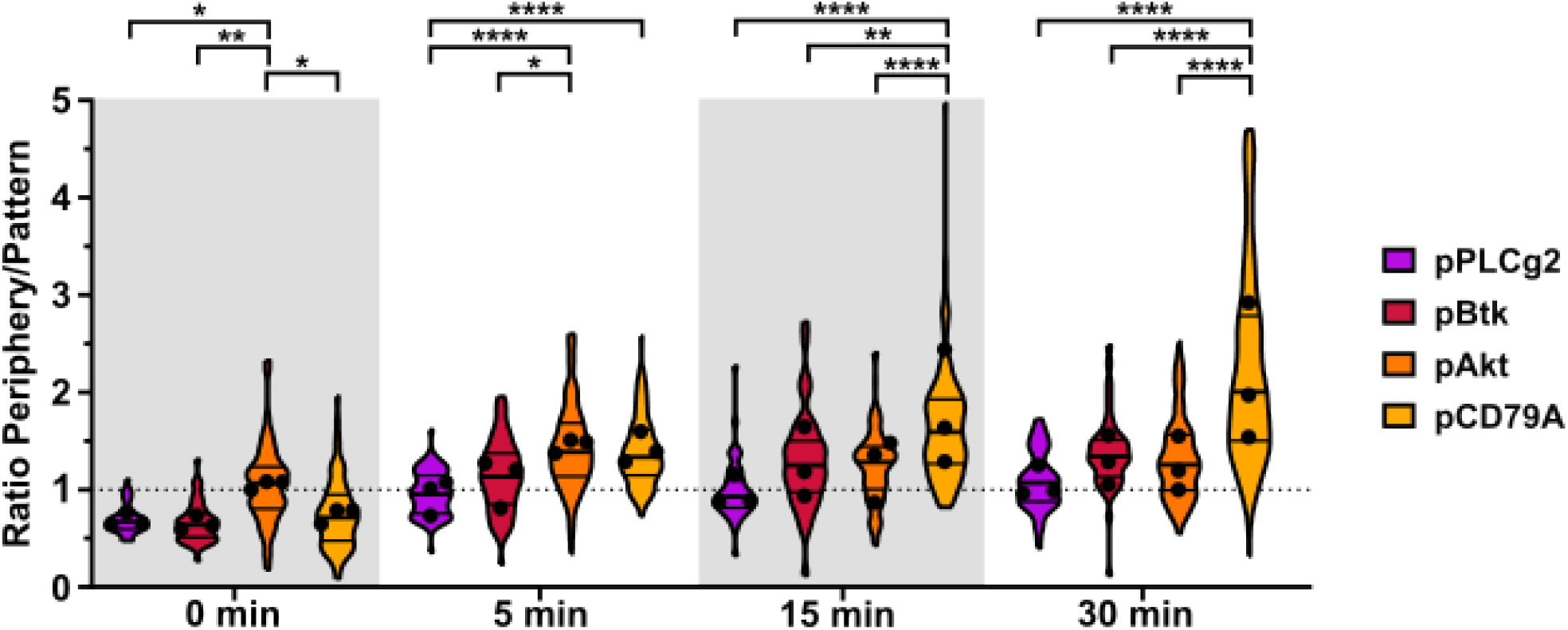
S4. Localization of BCR signaling proteins in the IS initiated from triangle micropatterns. Data from Figure 3 A-C, B cells activated on 15 µm triangle micropatterns, were analyzed for the ratio of periphery MFI/pattern MFI. Violin plot graphs with mean (central line), quartiles (top and bottom lines), and experiment means (dots). One-way ANOVA (n=3, 15-30 cells per condition per experiment).

**Figure S5:**
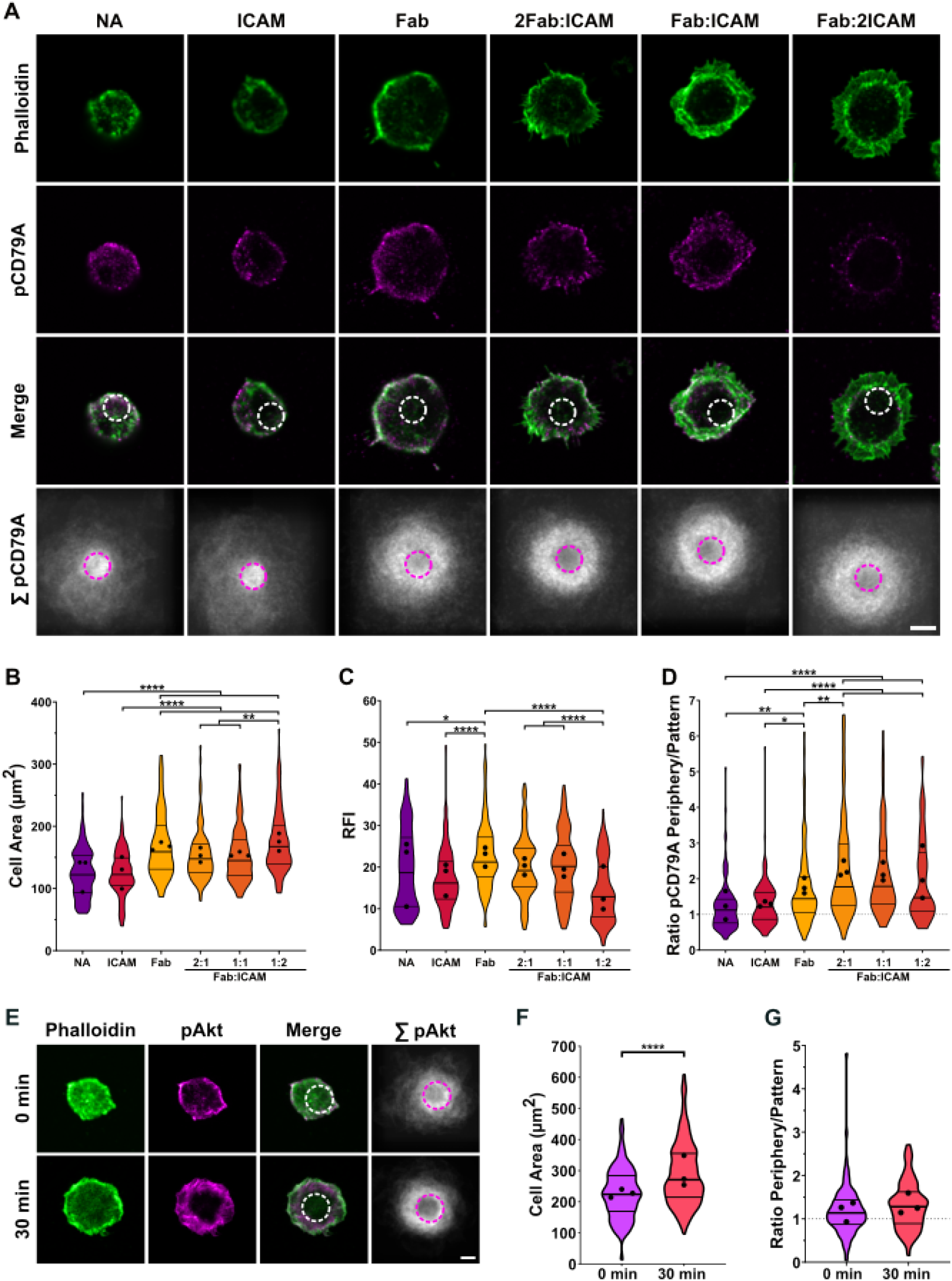
Addition of multiple ligands in dynamic micropatterns supporting B cell IS formation, and adaptation of dynamic micropatterns for T cell IS formation. (**A**) B cells were allowed to form immune synapses via seeding them on fibronectin/fibrinogen-coated circular 5 µm (dotted white or magenta lines) dynamic micropatterns and activating by the addition of different combinations of Strep-αIgM Fab and Strep-ICAM for 15 minutes. Samples were fixed and subjected for immunofluorescence analysis to visualize F-actin (green) and pCD79A (magenta). Representative images of single confocal microscopy slices are shown as well as sum images of phospho-proteins (white) of all cells per condition and staining. Scale bar 5µm. (**B-D**) Cell area (B), pCD79A mean intensity (C) and periphery MFI/pattern MFI ratio of pCD79A signal (D) from the data in (A) was analysed. Violin plot graphs with mean (central line), quartiles (top and bottom lines) and experiment means (dots) of the. One-way ANOVA (n=3, 30-50 cells per condition per experiment). (**E**)Jurkat T cells were seeded on fibronectin/fibrinogen-coated circular 10 µm (dotted white of magenta line) dynamic micropatterns for 20 minutes and activated by the addition of streptavidin-αCD3 and streptavidin-αCD28 to allow for the formation of the T cell IS. Samples were fixed and subjected for immunofluorescence analysis to visualize F-actin (green) and pAkt (magenta). Representative images of single confocal microscopy slices are shown as well as sum images of pAkt (white).(**F-G**)The data in (E) was analysed for the cell area (F), and ratio of phospho-Akt Periphery MFI/Pattern MFI (G). Violin plot graphs with mean (central line), quartiles (top and bottom lines) and experiment means (dots) are shown. One-way ANOVA (n=3, 30-50 cells per condition per experiment).

**Table S1:** List of antibodies used for immunofluorescence, western blotting and B cell activation.

| Antibody | Commercial house | Catalog number | Source species | Clone | Dilution |
| --- | --- | --- | --- | --- | --- |
| AF488 pPLCg2 (Y759) | BD Biosciences | 558507 | Mouse | K86-689.37 | 1:100 |
| AF647 pBtk (y223)/pIt(Y180) | BD Biosciences | 564846 | Mouse | N35-86 | 1:100 |
| Anti-phosphotyrosine | Merck | 05-321 | Mouse | 4G10 | 1:1000 |
| CD28 Monoclonal antibody | Invitrogen | 16-0289-81 | Mouse | CD28.2 |  |
| CD3 Monoclonal antibody | Invitrogen | 16-0037-85 | Mouse | OKT3 |  |
| Donkey anti-Goat IgG (H+L) Cross-Adsorbed Secondary Antibody, Alexa Fluor 488 | Thermo Fisher Scientific | A-11055 | Donkey | Polyclonal | 1:500 |
| Donkey anti-Rabbit IgG (H+L) AF647 | Invitrogen | A31573 | Donkey | Polyclonal | 1:500 |
| F(ab') Fragment Goat Anti-Mouse IgM, $\mu$ chain specific | Jackson ImmunoResearch | 115-006-020 | Goat | Polyclonal | |
| Fab Fragment Goat Anti-Mouse IgM, $\mu$ Chain Specific | Jackson ImmunoResearch | 115-007-020 | Goat | Polyclonal | |
| HRP Anti-beta actin antibody | Abcam | ab49900 | Mouse | AC-15 | 1:20000 |
| pAKT (Ser473) | Cell Signal Technology | 4058 | Rabbit | 193H12 | 1:100 |
| pCD79A (Tyr182) | Cell Signal Technology | 5173 | Rabbit | Polyclonal | 1:100 |

**Table S2:** List of reagents and consumables used for experimental work.

| Product | Commercial house | Catalog number |
| --- | --- | --- |
| 13 mm diameter coverslips | Epredia | CB00130RAC20MNZ0 |
| 25 mm diameter coverslips | Epredia | CB00250RAC20MNZ0 |
| ActiStain 488 Phalloidin | Cytoskeleton | PHDG1-A |
| Attofluor cell chamber for microscopy | Invitrogen | A7816 |
| CK666 | Sigma-Aldrich | 182515 |
| Fibronectin | Sigma Aldrich | F4759 |
| Fibronectin bovine plasma | Sigma-Aldrich | F4759 |
| Fluoromount G | Southern Biotech | 0100-01 |
| Fluoromount G with DAPI | Invitrogen | 00495952 |
| Glutaraldehyde | Sigma-Aldrich | G7776 |
| Immobilon Western Chemiluminescent HRP Substrate | Merck | WBKLS0500 |
| Latrunculin A | Merck Millipore | 428021 |
| Molecular Probes Fibrinogen From Human Plasma, Alexa Fluor 546 Conjugate | Thermo Fisher Scientific | F13192 |
| Paraformaldehyde | Sigma-Aldrich | 16005 |
| Para-nitro-Blebbistatin | Cayman Chemical | 24171 |
| Phalloidin AF488 | Cell Signalling Technology | 8878S |
| Phalloidin-iFluor 405 reagent | Abcam | ab176752 |
| PLL(20)-g[3.5]- PEG(2)/PEG(3.4)- biotin(50%) | SuSoS |  |
| Recombinant mouse ICAM-1-Fc Chimera (carrier-free) | BioLegend | 553004 |
| Streptavidin conjugation kit | Abcam | ab102921 |

